# Targeting metabolic adaptations in the breast cancer–liver metastatic niche using dietary approaches to improve endocrine therapy efficacy

**DOI:** 10.1101/2021.09.07.458711

**Authors:** Qianying Zuo, Ayca Nazli Mogol, Yu-Jeh Liu, Ashlie Santaliz Casiano, Christine Chien, Jenny Drnevich, Ozan Berk Imir, Eylem Kulkoyluoglu-Cotul, Nicole Hwajin Park, David J Shapiro, Ben Ho Park, Yvonne Ziegler, Benita S. Katzenellenbogen, Evelyn Aranda, John D. O’Neill, Akshara Singareeka Raghavendra, Debu Tripathy, Zeynep Madak Erdogan

## Abstract

Estrogen receptor-positive (ER^+^) metastatic tumors contribute to nearly 70% of breast cancer-related deaths. Most patients with ER^+^ metastatic breast cancer (MBC) undergo treatment with the estrogen receptor antagonist fulvestrant (Fulv) as standard-of-care. Yet, among such patients, metastasis in liver is associated with reduced overall survival compared to other metastasis sites. The factors underlying the reduced responsiveness of liver metastases to ER-targeting agents remain unknown, impeding the development of more effective treatment approaches to improve outcomes for patients with ER^+^ liver metastases. We therefore evaluated site-specific changes in MBC cells and determined the mechanisms through which the liver metastatic niche specifically influences ER^+^ tumor metabolism and drug resistance. We characterized ER activity of MBC cells both in vitro, using a novel system of tissue-specific extracellular matrix hydrogels representing the stroma of ER^+^ tumor metastatic sites (liver, lung and bone), and in vivo, in liver and lung metastasis mouse models. ER^+^ metastatic liver tumors and MBC cells grown in liver hydrogels displayed upregulated expression of glucose metabolism enzymes in response to Fulv. Furthermore, differential ERα activity, but not expression, was detected in liver hydrogels. In vivo, increased glucose metabolism led to increased glycogen deposition in liver metastatic tumors, while a fasting-mimicking diet increased efficacy of Fulv treatment to reduce the metastatic burden.

**Implications:** Our findings identify a novel mechanism of endocrine resistance driven by the liver tumor microenvironment. These results may guide the development of dietary strategies to circumvent drug resistance in liver metastasis, with potential applicability in other metastatic diseases.

## Introduction

About one-third of women with early-diagnosed non-metastatic breast cancer later develop metastatic disease (1), and 90% of breast cancer deaths result from metastases that is intractable to treatment (2,3). Among US women with breast cancer, an estimated 168,000 had metastatic breast cancer (MBC) in 2020 (4). This condition confers poor outcomes, with a median survival of approximately 18–24 months from metastatic tumor diagnosis (5). These adverse outcomes underscore the need to identify mechanisms promoting metastatic disease and treatment resistance.

MBC commonly spreads to the liver (6,7). Importantly, patients with liver metastasis (40.8%) have significantly increased death risk, similar to brain metastasis, and a disproportionally higher mortality rate compared to lung (36.8 %) or bone metastases (67%) (8-11). About 70% of metastatic tumors express estrogen receptor alpha (ERα), rendering MBC responsive to endocrine-based therapies such as Fulvestrant (Fulv) (12,13). This ERα antagonist is the only clinically approved selective ER degrader prescribed either alone or in combination with CDK4/6 or PI3K inhibitors (for PIK3CA mutant tumors) to treat MBC, independent of metastatic site. Notably, patients with liver metastases have poorer response to Fulv compared to patients with bone or lung metastases (6,7,14). Left untreated, survival for patients with MBC in liver is typically 4–8 months (15). For those receiving endocrine/biotherapy or chemotherapy, median times to progression are in the range of 20–30 months with initial therapy, and these times shorten progressively with subsequent treatments until clinical resistance develops (10). Therefore, there is a critical need for novel therapeutic approaches that will provide a durable therapy response or cure for patients with ER^+^ liver MBC.

Metastatic tumor phenotypes and treatment responses result from complex interactions between tumor cells and components of the tumor microenvironment, including inflammatory cells, fibroblasts, and biochemical composition and physical properties of the extracellular matrix (16). A crucial aspect of this microenvironment is the metabolic state. Cancer cells exhibit a remarkable metabolic plasticity (17,18), whereby they adapt to new metastatic environments by rewiring their metabolic pathways (16). Yet, these changes not only provide building blocks to sustain the cells’ biological functions, they also create novel vulnerabilities that are potentially exploitable through pharmacological or dietary interventions.

We recently showed that MBC cells grown in tissue-specific extracellular matrix hydrogels to tune their microenvironment displayed altered metabolic profiles that rendered the cells vulnerable to metabolic inhibitors; notably, treatment with these inhibitors improved MBC cell responses to endocrine therapy (19). Previous work using syngeneic mouse models showed metastatic site-specific metabolic adaptations in ER-negative breast tumors (20,21). Yet, the impact of the liver microenvironment on ER^+^ MBC cell metabolic reprogramming and response to ER-targeting agents is unknown.

Here, we sought to identify how the metabolism–cancer nexus in liver affects the response of metastatic ER^+^ tumors to Fulv, focusing particularly on tumor-intrinsic metabolic mechanisms arising following Fulv exposure. Spatial and bulk transcriptomic, and metabolic analysis of MBC cells and in vivo xenograft tumors identified an upregulation of glucose metabolism in the liver niche in response to Fulv. We fed mice with diets with differing carbohydrate levels to modulate metastatic burden and improve Fulv responses. Our findings delineating the underlying causes of the unique MBC response to Fulv in the liver should inform the design of future clinical trials to test efficacy of combining dietary interventions with endocrine therapies to improve treatment response.

## Materials and Methods

### Clinical Data and Analysis

The breast cancer management database at The University of Texas MD Anderson Cancer Center was used to identify patients for the current study, with the following inclusion criteria: age ≥ 18 years, diagnosed with de novo or recurrent ER^+^/HER2^-^ MBC between 1997 and 2021, with metastasis to the liver and receipt of Fulv in their metastatic setting. Study data were collected and analyzed with approval from the Institutional Review Board (IRB) at the University of Texas MDACC. A waiver of consent was obtained due to the retrospective nature and minimal patient risk. To evaluate the difference in overall mortality among patients with MBC to the liver, we analyzed 3388 patients with MBC in a MD Anderson Cancer Center cohort study. Patients with either primary or secondary liver metastasis were identified as having hepatic involvement. All other patients with MBC were identified as having non-liver metastasis. Kaplan-Meier methods were used to visualize overall survival and survival after metastasis. The Cox proportional hazards model was then used to compare the survival between patients with metastasis to the liver versus metastasis to other sites. These models were adjusted for age at diagnosis, BMI, race, and tumor stage.

### Cell Culture

All cell lines were obtained from American Type Culture Collection (Manassas, VA) unless indicated otherwise. MCF7 (ATCC HTB-22) (RRID:CVCL_0031) and T47D (ATCC HTB-133) (RRID:CVCL_0553) parental cells were cultured in RPMI-1640 medium with NEAA salts (Sigma, St Louis, MO, USA), 5% fetal bovine serum (FBS) (HyClone, Logan, UT), 100 μg/mL penicillin/streptomycin (Invitrogen, Carlsbad, CA, USA) and 50 mg/mL Gentamicin (Gibco, Gaithersburg, MD). MCF7-ESR1^D537S^ and - ESR1^Y537S^ cells (RRID:CVCL_0031) were generated as described in (22) and were cultured in Dulbecco’s Modified Eagle Medium (DMEM) with NEAA salts, 10% FBS, 100 μg/mL penicillin/streptomycin, and 50 mg/mL gentamicin. T47D*-*ESR1^D537G^ and T47D-ESR1^Y537S^ cells (RRID: CVCL_0553) were cultured in Modified Eagle Medium (MEM) with NEAA salts, 10% FBS, 100 μg/mL penicillin/streptomycin, and 50 mg/mL gentamicin. Cell line authentication was performed by checking activity and expression of ERα, proliferative responses to ER agonists and antagonists for all cell lines, and sequencing of MCF7-ESR1^Y537S^ and MCF7-ESR1^D537G^ as described (23). Cell lines were monitored for Mycoplasma contamination regularly using Mycoplasma Detection Kit (VWR, 89510-164).

### 3-D Cell Culture Models

IN SITE Metastasis Kit (Xylyx Bio, Inc., NY) containing TissueSpec Bone (MTSBN101), Liver (MTSLV101), and Lung (MTSLG101) ECM Hydrogels, were used to model tumor microenvironments according to the manufacturer’s protocol. Briefly, 2×10^3^ cells were encapsulated in corresponding extracellular matrix (ECM) hydrogels by mixing them with tissue culture matrix. A mixture volume of 100 uL/well was placed in 96-well plates in triplicate. Plates were incubated at 37°C in a humidified environment with 5% CO_2_ for at least 45 minutes to achieve gelation. Cells were treated with media containing Veh or 1μM Fulv every Monday and Friday for three weeks. Oncosphere formations were visualized by Invitrogen EVOS XL Core Light Microscope (4X and 25X magnifications) (Waltham, MA, USA). OpenCFU colony counting software (http://opencfu.sourceforge.net/) was used to automatically count colony number and size. For Fulv dose response experiments MCF7-ESR1^Y537S^ cells were seeded on polymerized hydrogels at a density of 3.0 × 10^3^ cells/well (96 well plate) followed by addition of culture media (DMEM) at Day 0 for 24 hours. After 24 hours Fulv treatment was started using the following concentrations: 0.001, 0.01, 0.1, 1, 10μM. Final concentration of 0.01% Ethanol was used as control (veh) treatment. Experiments were conducted in triplicates. Plate was kept for 10 days, with treatments performed on Days 1, 4, and 7. Images were taken on day 10 using Cytation 5 Imaging Multi-Mode Reader (Biotek®). Images were further analyze using ImageJ. (24) Images were imported and threshold adjusted. Analyze particles tool was used to count colonies and measure colony area. Ordinary one-way ANOVA test was performed to compare the means of the unmatched concentrations. No matching or pairing was selected and multiple comparisons were made: the mean of each dose was compared with the mean of control dose (Veh). All statistical analyses were completed by using GraphPad Prism 8 software (RRID:SCR_002798).

### Western Blot

Western blot analysis used specific antibodies against β-actin (Sigma Aldrich, St. Louis) and ERα (F10, Santa Cruz Biotechnology, Santa Cruz, CA). MCF-7 cells were seeded at 2–4×10^5^ cells in 10-cm dishes in 5 mL growth media. The next day, cells were treated with fresh media and collected 15 min later into 250 μL lysis buffers. Cell lysate was prepared using RIPA buffer. Samples were sonicated three times for 10 s to shear the DNA. Ten micrograms of protein were loaded onto 10% SDS gels. ERα primary antibody was used at 1:500 except for β-actin (1:5,000). Secondary antibodies are from Licor biosciences (Goat anti rabbit IRDdye 800 and goat anti mouse IRDye 680). Proteins were visualized using Odyssey LI-COR Imaging System that detect infrared dye on the secondary antibodies.

### In Vivo Xenograft Study

Four-week-old, ovariectomized, NOD SCID gamma (NSG) immunodeficient female mice were obtained from The Jackson Laboratory (Bar Harbor, ME). All experiments involving animals were conducted in accordance with National Institutes of Health standards for the use and care of animals, with protocols approved by the University of Illinois at Urbana-Champaign (UIUC; IACUC protocol #20158). After one week of acclimatization to the housing facility, 1×10^6^ MCF7-ESR1^Y537S^ or MCF7-ESR1^D538G^ cells resuspended in 1% PBS were injected via tail vein and randomized animals to indicated treatment groups. Mice were housed in 12-h light-dark cycle.

#### Diet and metastatic burden study

Three different diets were implemented for this study: Control (F4031, Bioserv, Flemington, NJ), High-fat Diet (HFD) (F3282, Bioserv), and Fasting-mimicking Diet (FMD) (F3666, Bioserv). We randomized N=8 mice to one of the two treatments Vehicle (Veh) or Fulv in each diet group. Fulv (Sigma) was dissolved in 10% DMSO and 90% corn oil and administrated via intramuscular injection (100 mg/kg) twice a week (Monday, Friday) for four weeks.

#### FMD and Fulv response study

In our diet studies using MCF7-ESR1^Y537S^ cells, metastatic tumor bioluminescence radiance had an average of 1.9×10^6^ p/sec/cm^2^/sr and standard deviation of 1.5×10^6^ p/sec/cm^2^/sr. Based on these values and Type 1 error of 5% and Type II error of 5%, to observe a significant tumor response, 8 mice per group were randomized to the Fulv treatment or control groups (i.e., Control/Veh; Control/Fulv; FMD/Veh; FMD/Fulv). Fulv was dissolved in 10% DMSO and 90% corn oil and administrated via intramuscular injection (100 mg/kg) twice a week (Monday, Friday) for four weeks.

For all experiments, food intake and body weight were measured twice weekly for the study duration. After six weeks of treatment, mice were euthanized, and organs were harvested. Tumor growth was monitored over time using an in vivo bioluminescence imaging (IVIS) system (PerkinElmer). Tumor burden was assessed by counting the final metastatic tumor nodules at necropsy. Aliquots of the samples were either flash frozen, frozen in RNAlater (Thermo Scientific, Waltham, MA, USA) for RNA isolation, or crosslinked in 10% neutral buffered formalin at room temperature (Millipore Sigma, Burlington, MA) for histological analysis.

### Histology

Tissues were sectioned (5 micron thick) using a microtome (Leica RM1255, Austria). For ERα immunostaining, tissues were first. Deparaffinized. Then, they were hydrated through graded alcohols to water. A citrate buffer at pH 6.0 was applied in a steamer for 1 hour for the antigen retrieval. Samples were blocked in hydrogen peroxide for 10 min followed by Background buster (Innovex Biosciences, Richmond, CA) incubation for 10 minutes to remove non-specific protein staining. After rinsing samples with TBS-Tween solution, pH 7.6, samples were incubated with anti-ERα (F10, Santa Cruz Biotech) (RRID: AB_631470) primary antibody overnight at 4° C. After rinsing with TBS-Tween solution at pH 7.6, samples were stained with secondary anti-rabbit and anti-mouse HRP-Polymer (Biocare Medical, Concord, CA) for 30 min. Finally, after DAB (Innovex, Richmond, CA) incubation for 5 min and counterstaining with hematoxylin, samples were dehydrated, and mounted on slides. Nanozoomer Slide Scanner (Hamamatsu, Japan) was used to visualize the samples at 80X magnification, and positive staining quantification was performed using NDP software.

For the Periodic Acid Schiff’s (PAS) staining, paraffin sections were deparaffinized in xylene and rehydrated through graded alcohols to water. Then they were placed in 0.5 % periotic acid for ten minutes. After rinsing well in water, they were placed in McManus Schiff’s Reagent (Newcomer Supply, Inc., Middleton, Wisconsin) for 10 minutes. Following a 30 second rinse in 0.55 % Potassium Metabisulfite, the slides were placed in warm running tap water for10 minutes. They were then counterstained with modified Harris hematoxylin (Thermo Fisher Scientific, Kalamazoo, MI). Slides were then dehydrated, cleared in xylene and mounted. For glycogen digestion, following deparaffinization, the slides were placed in 1% amylase in a 37 °C incubator for 30 minutes before proceeding with the PAS stain.

### Gene Expression Analysis

To identify gene sets regulated in different ECMs, *S*elective *E*strogen *R*eceptor *M*odulators (SERM), or by Fulv in liver metastatic tumors, RNASeq analysis was performed. For ECM experiments, 30-mm cell culture plates were coated with Native Coat ECMs for liver, lung, or bone (Xylyx). Then, these plates were incubated at 37ºC in a humidified environment with 5% CO_2_ for at least for 1 hour. MCF7-ESR1^Y537S^ cells were seeded on coated plates after removing native coat mixtures at a density of 2×10^3^ cells/well in DMEM. Cell lysates were collected using Trizol after 24 h of culture. MCF7-ESR1^Y537S^ cells were treated with Vehicle (Veh, 0.5% EtOH), 10^−6^ M 4-hydroxytamoxifen (4OHT) (Sigma), 10^−6^ M Fulv (Sigma), or 10^−6^ M Palbociclib (Palb) (Sigma) for 24 h. Concentrations of drugs were selected based on our previously published studies (19,25) and clinical data (26-28). Experiments were performed in triplicate. For cell line xenograft tumors of MCF7-ESR1^Y537S^ cells, Veh- or Fulv-treated tumors were homogenized in 1 mL of TRIzol reagent (Life Technologies, Carlsbad, CA, USA). Total RNA was extracted with TRIzol reagent according to the manufacturer’s protocol and cleaned using a kit (QIAGEN, Hilden, Germany). RNA quality was assessed using bioanalyzer. Total RNA from each sample (three per treatment group) was sequenced at the UIUC sequencing center, and data were generated in Fastqc file format to compare transcript abundance between the four treatment groups.

#### Preprocessing and quality control

Trimmomatic (Version 0.38) was used to trim Fastqc files containing raw RNA sequencing data (RRID:SCR_011848) (29). After the reads were mapped to the human reference genome (GRCh37) or mouse reference genome (GRCm38) from the Ensembl (RRID:SCR_002344) (30) database and STAR tool was used for alignment (RRID:SCR_015899) (Version 2.7.0f) (31). SUBREAD (Version 1.6.3) was used to generate read counts (RRID:SCR_009803) (32), and feature counts were exported for statistical analysis in R. R package edgeR (Version 3.24.3) (RRID:SCR_012802) was used to perform quality control and normalization (33). R limma package (Version 3.38.3) was used for Empirical Bayesian statistical analysis on the fitted model of the contrast matrix (RRID:SCR_010943) (34,35). We considered genes with fold-change >2 and p-value <0.05 (Benjamini and Hochberg multiple test correction for each gene, for each treatment) as statistically significant, differentially expressed. Cluster3 software was used for clustering the differentially expressed genes, which was visualized using Treeview Java (RRID:SCR_016916). Principal components analysis (PCA) was performed using StrandNGS (Version 3.1.1). To identify GO terms, and gene sets associated with different conditions Gene set enrichment analysis (GSEA, RRID:SCR_003199) (36,37) was used.

Visium Spatial transcriptomics libraries (10X Genomics) were constructed at the Carl R. Woese Institute for Genomic Biology Core Facility and the DNA Services laboratory of the Carver Biotechnology Center at the University of Illinois at Urbana-Champaign. Briefly, frozen tissues were sectioned and placed onto a Tissue Optimization Slide with eight capture areas, each having thousands of spots with poly-dT capture probes. Tissue sections were permeabilized across a time-course from 3 to 30 minutes. The mRNAs from the tissue anneal to the oligos, were converted to cDNA by reverse transcription with a fluorophore and the sections were evaluated with the Zeiss Axio Observer Z1 microscope. The permeabilization condition producing the brightest image with the least diffusion were chosen for the processing of the tissues. A permeabilization time of 12 minutes was selected for these samples. Next, tissue sections were placed onto the Visium Spatial Gene Expression slide, which contains 4 capture area squares 6.5mm x 6.5mm. Each square has approximately 5,000 spots with barcoded poly-dT oligos. Tissue sections were fixed, stained with hematoxylin and eosin (H&E) and visualized with the Hananatsu Nanozoomer microscope. After permeabilization, messenger RNAs were converted into spatially-barcoded cDNAs following the 10x Genomics protocol. The double-stranded-barcoded cDNAs are then denatured and converted into a sequencing-ready, dual-indexed libraries. The final libraries were quantitated on Qubit and the average size determined on the AATI Fragment Analyzer (Advanced Analytics, Ames, IA), then diluted to 5nM concentration and further quantitated by qPCR on a Bio-Rad CFX Connect Real-Time System (Bio-Rad Laboratories, Inc. CA). The libraries were pooled by qPCR value and capture spot coverage and sequenced on an Illumina NovaSeq 6000 to a length of 28nt (read 1, contains the spot barcode and unique molecular identifier used for removing PCR duplicates), 10nt for each index (libraries contain unique dual indexes to prevent index switching) and 150nt for read 2 (the cDNA read) to a minimum depth of at least 100,000 cDNA read 2 per spot. Fastq.gz files were generated and demultiplexed with SpaceRanger 1.3.0. Data was processed and visualized using Space Ranger Analysis Pipelines and Loupe browser using a combined human (GRCh38) and mouse (GRCm39) reference based on Ensembl Release 104. Visium uses the cDNA barcodes to associate the transcripts to an X-Y coordinate on the slide, which can then be used to overlay the H&E-stained image with the transcript information from a spatial viewpoint. The R package Seurat was used to read in both the count data and the images. To identify spots that were primarily mouse host, we calculated the proportion of all UMIs that were from human genes and excluded spots that were <= 25% human. All mouse genes were also removed and then all 4 samples were normalized together using sctransform, then principal components analysis was performed and the top 30 PCs were used in both nearest neighbor cluster calling and UMAP dimension reduction.

### Metabolomics Analysis

MCF7-ESR1^Y537S^ cells were seeded at a density of 2×10^5^ cells/plate as explained above. After 24 hours, acetonitrile/isopropanol/water extraction was performed. Metabolite extracts were stored at -80ºC until submitted to the Metabolomics Center in the Roy J Carver Biotechnology Center at UIUC. To detect and quantify the metabolites gas chromatography-mass spectrometry (GC/MS) analysis was performed. We detected the metabolite profiles using an Agilent GC-MS system (Agilent 7890 gas chromatograph, an Agilent 5975 MSD, and an HP 7683B autosampler). Spectra of all chromatogram peaks were evaluated using AMDIS 2.71 and a custom-built database with 460 unique metabolites. All known artificial peaks were identified and removed before data mining. All data were normalized to the internal standard in each chromatogram to ensure comparison between samples. This analysis identified about 200 metabolites. We used web-based Statistical Analysis tool of MetaboAnalyst software (RRID:SCR_015539) version 4.0 (38). Features with more than 50% missing data points were removed. Data were log transformed and scaled using the auto-scaling feature. A clustered heatmap of class averages of top 25 metabolites was generated.

To visualize the separation between No ECM and Liver ECM groups’ metabolic profiles VIP scores for the top 25 metabolites that discriminated between treatment groups were calculated and displayed using the Partial least squares discriminant analysis (PLS-DA) tool. Enrichment analysis and pathway analysis were used to identify metabolic pathways associated with enriched metabolites (fold change >2 or <0.05) as previously described (39-42).

### Seahorse Metabolic Profiling Assays

Seahorse XFp plates were coated with Native Coat ECMs for bone, lung, or liver (Xylyx), and were incubated at 37°C in a humidified environment with 5% CO_2_ for at least for 1 hour. MCF7-ESR1^Y537S^ cells were seeded at a density of 3×10^4^ in corresponding treatment media without phenol red in each well of the XFp Cell Culture miniplates (Seahorse Bioscience Inc., Billerica, MA, USA). The next day, the cartridges were hydrated with the calibration solution and were kept overnight in a non-CO_2_ incubator at 37°C. To ensure ECAR and OCR values were normalized to the cell number, we monitored the cell number changes after 24 h of treatments. On the assay day, XF Base Media without phenol red (Seahorse Bioscience Inc., Santa Clara, CA) supplemented with 10 mM L-glucose, 2 mM L-glutamine (Gibco), and 1 mM sodium pyruvate (Gibco, Waltham, MA) was used to wash the cells. ECAR (mpH/min) and OCR (pmol/min) values were obtained using Seahorse XFp Cell Energy Phenotype Test Kit (Seahorse Bioscience Inc.). Experiments were performed in triplicate and repeated at least three times.

### ChIP-seq Analysis

ChIP-seq analysis was performed as described previously using MCF7-ESR1^Y537S^ cells (43,44). ERα–DNA or IgG–DNA complexes were immunoprecipitated using ERα-specific F10 and HC20 antibodies (Santa Cruz Biotech, 3:100 dilution). ChIP DNA was obtained from three pooled biological replicates. Libraries were prepared according to Illumina Solexa ChIP-Seq sample processing (San Diego, CA), and single-read sequencing was performed using the Illumina HiSeq 2000. Sequences generated were mapped uniquely onto the human genome (hg18) by Bowtie2 (RRID:SCR_016368). The MACS (model-based analysis of ChIP-seq) algorithm was used to identify enriched peak regions (default settings) with a P value cutoff of 6.0e−7 and FDR of 0.01, as we have described (41,45). BETA method (RRID:SCR_005396) was used to map ERα binding sites to regulated genes in different ECMs relative to 2D. Bed files with high-confidence binding sites from MACS and differentially expressed gene lists from ECM gene expression dataset were used. The binding sites that are located within 100 Kb of a differentially regulated gene (FDR<1%) are evaluated based on regulatory potential method to do the control region prediction as described (46).

### Statistical Analyses

Data from all studies were analyzed using a one-way analysis of variance (ANOVA) to compare different ligand or ECM effects, or a two-way-ANOVA model to compare time-dependent changes. Unpaired t-tests with a Bonferroni correction was used to analyze normally distributed data and to identify treatments that produced significantly different results (* p <0.05, ** p <0.01, *** p <0.001, **** p <0.0001). Data that were not normally distributed were analyzed using Mann-Whitney test for nonparametric data (* p <0.05, ** p <0.01, *** p <0.0001). Statistical significance was calculated using GraphPad Prism 9 for Windows.

### Data Availability

Gene expression data were submitted to the GEO database (GSE184156) and will be available from the day of the acceptance of the manuscript.

## Results

### Patients with liver metastatic breast tumors respond poorly to Fulv

Small cohort studies reported that patients with liver metastases are less responsive to Fulv compared to patients with bone or lung metastases (6,7,14). To validate these findings in a larger cohort, we analyzed data from an ongoing trial for patients with ER^+^ MBC. We identified 1556 (46%) patients in our cohort with liver metastasis. Median overall survival in patients with liver metastasis was 10 years (9.6, 10.4), which was significantly shorter than those with metastasis elsewhere. The hazard ratio was 1.46 (1.34, 1.59), indicating an increased risk of mortality associated with liver metastasis across all treatment regimens (**Fig.1A**). Those with liver metastasis also had worse survival post-metastasis, with a median survival of 5.8 years (5.6, 6.0) compared to 6.7 years (6.5, 7.0) in the non-liver metastasis cohort (**Fig.1B**). We also analyzed the impact of chemotherapy on survival in patients who were treated with fulvestrant therapy. 1726 (58%) patients received chemotherapy before they present with metastatic disease **(Figure S1A and S1B)**. 1467 (49.4%) patients received chemotherapy after metastasis **(Figure S1C and S1D)**. While chemotherapy before metastasis improved survival of patients with liver metastasis (Figure S1A and S1B), chemotherapy after metastasis did not improve survival of patients with liver metastasis (Figure S1C and S1D). Out of 1414 patients in the cohort with liver metastasis, 197 (14%) had liver as the first site of metastasis. Patients with first metastasis to the liver did not significantly differ in overall survival or survival after metastasis when compared to those who developed liver metastasis later (**Figure S1E and S1F**).

**Figure 1:**
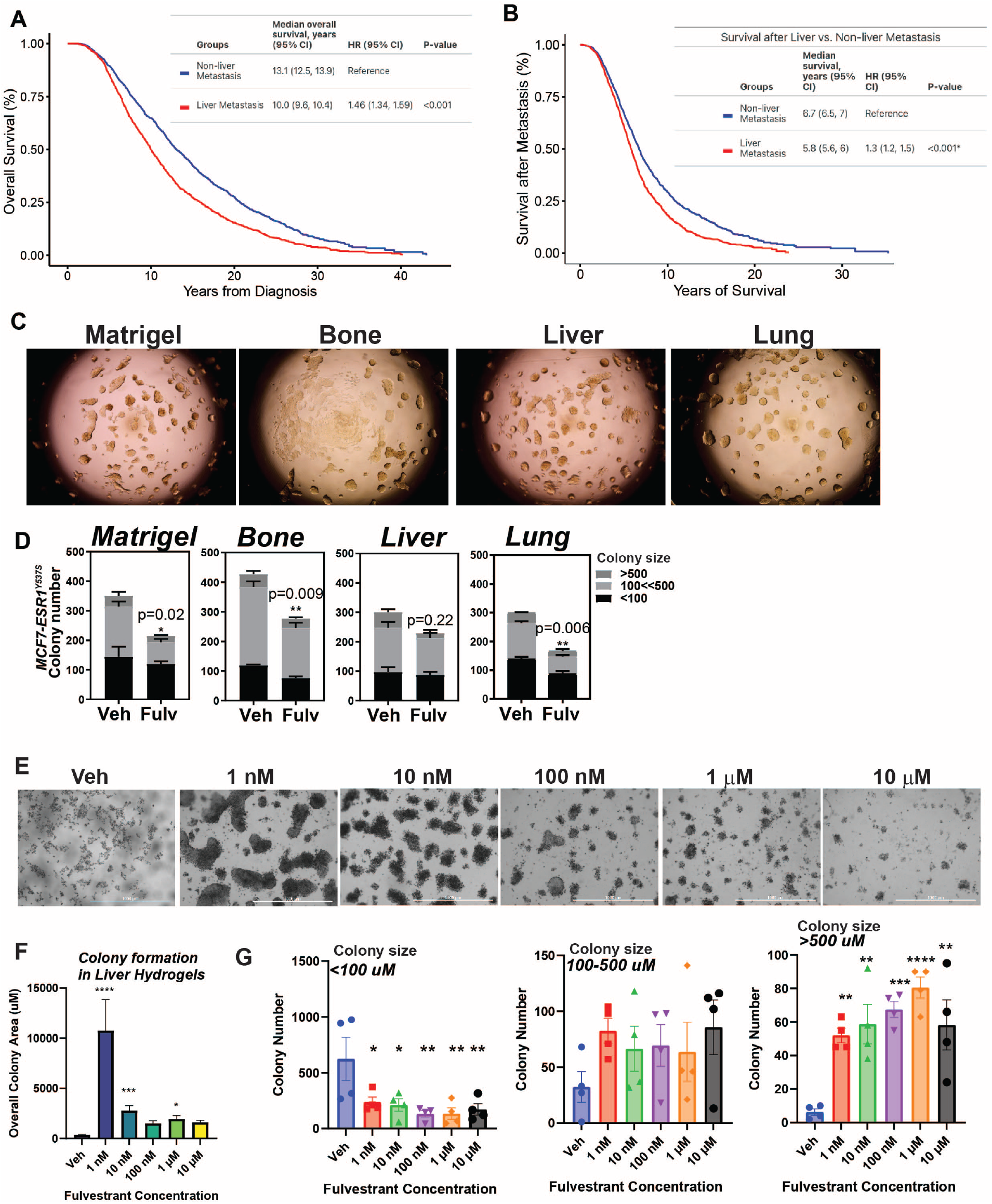
Patients with liver metastatic breast tumors experience poor response to Fulv. MD Anderson Cancer Center cohort study involved 1832 patients without liver MBC and 1556 with liver MBC. Number of dead and alive patients with and without liver MBC were compared using Fisher’s exact test. Patients with ER^+^/HER2^-^ liver metastatic tumors have shorter overall survival (**A**) and shorter survival after metastasis (**B**). Log-rank test was used to compare survival curves. **C)** Colony formation assay for MCF7-ESR1^Y537S^ cells in Matrigel or decellularized hydrogels from different metastatic tissues and **D)** and efficacy of 1 μM Fulv to reduce colony number. Colonies were treated for 3 weeks. Unpaired t-test, P-values are indicated. **E)** Fulv dose response for colony formation of MCF7-ESR1^Y537S^ cells in liver hydrogels. **F)** Quantification of colony size and number from Fig. 1E. **G)** Quantification of colony number for colonies in different area brackets.

Estrogen receptor alpha gene (ESR1) mutations were identified in 15-40% of patients with ER^+^ metastatic tumors (47-50). Since these ESR1-activating mutations within the ligand-binding domain of ERα are enriched only in metastatic tumors (47-50), particularly in visceral tissue metastasis including liver metastases (51-53), we analyzed enrichment of ESR1 mutations in liver metastatic tumors using two publicly available datasets (54,55). This analysis confirmed enrichment of ESR1 mutations in liver metastatic tumors (**Figure S1G and S1H**). In our cohort, among 2971 patients treated with fulvestrant, estrogen receptor 1 (ESR1) gene mutations were observed in a minority (127, 4%). Of those who had ESR1 mutations, 62 (49%) had metastasis to the liver. A lower prevalence of liver metastasis was found in patients without *ESR1* mutations (45%). In patients without ESR1 mutations, metastasis to the liver was associated with worse overall survival (HR 1.47; CI 1.35-1.6; P<.001) compared to metastasis to other sites (**Figure S1I**). In ESR1 mutation group, patients with liver metastasis had a shorter median overall survival of 19.25 months than those with non-liver metastasis who had median overall survival of 27.97 months (HR 1.63, P<0.1) (**Figure S1J**).

To characterize ER antagonist responses of MBC cells and test potential efficacy of combination therapies, we previously used decellularized hydrogels from different metastatic tissues (19). To examine the differences in Fulv response when MBC cells are grown in different hydrogels, we used MCF7-ESR1^Y537S^ cells (**Fig. 1C**). Consistent with the clinical observation, breast cancer cells grown on liver hydrogels were less responsive to Fulv compared to cells grown on other hydrogels (**Fig. 1D, and Figure S2**). Fulv increased colony area and number of colonies in a dose dependent manner, particularly for bigger colonies (**Figure 1E, 1F and 1G**).

### ER^+^ MBC cells display distinct transcriptomes and ER cistromes in different tissue-specific ECMs

To determine the molecular changes associated with different metastatic environments, we compared gene expression profiles from MCF7-ESR1^Y537S^ cells grown on plastic (2D), bone, liver, or lung ECM. MBC cells grown on different ECMs exhibited distinct gene expression profiles (**Fig. 2A and 2B**). Despite a lack of change in ERα protein expression (**Fig. 2C**), upregulation of gene sets associated with classical ERα signaling were less common in MBC cells grown in liver ECM compared to bone or lung ECMs (**Fig. 2D**). To investigate why classical ERα target genes were downregulated on liver ECM, we performed ChIP-seq analysis, which revealed that ERα recruitment to chromatin was significantly altered in a niche-specific manner (**Fig. 2E**). Overall, there were more ER binding sites in cells grown on bone ECM compared to lung or liver (**Fig. 2F**). These binding sites mapped to classical ER target genes. Estrogen response element was the most enriched motif in ER binding sites, suggesting a direct ERα binding to chromatin rather than tethering (**Table S1**).The fraction of distal intergenic binding sites was increased in cells grown on liver ECMs compared to other sites (**Fig. 2G**). Consistent with **Fig. 2D**, percent of upregulated genes with binding sites was lowest for liver (**Fig. 2H**). ER binding sites in cluster 2 (**Fig. 2E, C2**), present in cells grown in 2D plastic or bone ECMs, were lost or reduced in cells grown on liver or lung ECM (**Fig. 2I)**. Conversely, a new cluster of binding sites, cluster 3 (**Fig. 2E, C3**), was present only in cells grown in liver ECM, indicating the liver metastatic niche can promote a distinct pattern of ERα recruitment (**Figure 2J, Figure S2C, S2D and S2E**). These binding sites mapped to genes involved in metabolic regulation and insulin signaling, suggesting that these newly gained sites might be responsible for an altered metabolic phenotype.

**Figure 2:**
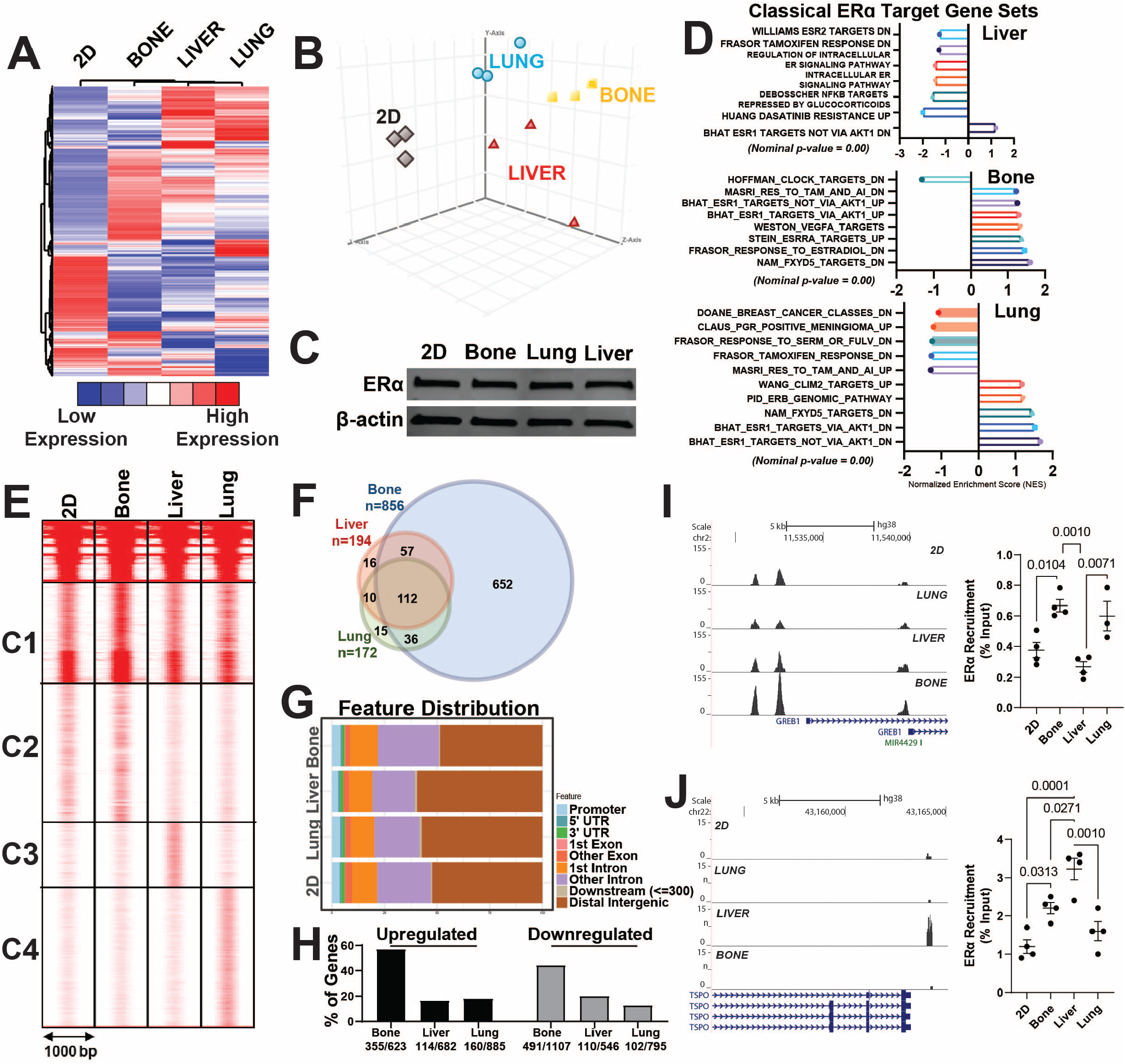
ER^+^ MBC cells display distinct transcriptomes and ER cistromes when grown in different ECMs. **A)** Hierarchical clustering of RNASeq data of MCF7-ESR1^Y537S^ cells grown on 2D (plastic), bone, liver, or lung Native Coat ECMs for 24 hours. Total RNA was isolated, and sequencing was performed using three samples from each treatment group. Differentially expressed genes were determined with P <0.05 and expression fold change >2. **B)** Principal component analysis of gene expression data using differentially expressed genes list from **A**, which shows distinct gene expression profiles associated with different ECMs. **C)** ERα protein expression was examined using western blotting. β-actin was used as a loading control. **D)** Gene-set enrichment analysis of gene sets that were enriched in related genes as classical targets of ERα action dataset. **E)** ERα ChIP-Seq in MCF7-ESR1^Y537S^ cells grown on plastic (2D), bone, liver, or lung Native Coat ECMs. ERα–DNA complexes were pulled down using ERα antibodies. Three biological replicates were pooled and sequenced. Clustering of ERα-binding sites in ECMs was done using seqMINER software. The ERα-binding sites were separated into four clusters of characteristic patterns: C1, C2, C3, and C4. **F)** Venn diagram analysis of ER binding sites in different tissue-specific ECMs **G)** Distribution of bindings sites relative to transcription start site **H)** Percentage of Metastatic site specific ERα binding sites that map to up- or downregulated genes in each metastatic site respectively. **I)** Example histogram and Q-PCR showing ERα recruitment to GREB1 binding sites in cells grown on different Native Coat ECMs. **J)** Example histogram and Q-PCR showing ERα recruitment to TSOP binding sites in cells grown in different ECM.

### Glucose dependence of MBC cells increases in liver ECM

To assess the impact of metastatic ECM on MBC metabolism, we grew MCF7-ESR1^Y537S^ cells on different ECMs. Growth of these cells on liver ECM caused an increase in both the glycolysis and oxidative respiration in Seahorse cell phenotype tests (**Fig. 3A**). To identify specific pathways impacted by liver ECM, we performed a metabolomic analysis. Growth of MBC cells on different ECMs resulted in distinct metabolite profiles when a PLS-DA classification was performed (**Fig. 3B**). While the abundances of fatty acid and glucose metabolism pathway metabolites were increased on liver ECM, amino acid and nucleotide metabolism-related metabolites were increased on lung ECM (**Fig. 3C**). To dissect the specific fuel dependency on different ECMs, we assayed glycolytic and oxidative respiration, which showed that liver ECM but not ECM from bone or lung increased glycolytic respiration in the presence of full media, or media with glucose or pyruvate compared to cells that were grown on plastic (2D) (**Fig. 3D**). Liver ECM led to an increase in oxidative respiration only when cells were supplemented with full media (**Fig. 3E**). A Seahorse cell phenotype test in different media showed that MBC cells grown in media with glucose had an increase in glycolytic potential when also grown on liver ECM (**Fig. 3F**). Finally, we performed metabolomic profiling in MBC cells grown on liver ECM that were treated with Veh or Fulv. This analysis further showed that relative abundance of glucose metabolism-associated metabolites including pyruvate were increased in response to Fulv (**Fig. 3G**), however impact of the Fulv treatment on OCR and ECAR values were minimal, either not changing or slightly increasing OCR in different breast cancer cell lines in liver ECM (**Figure S3**).

**Figure 3:**
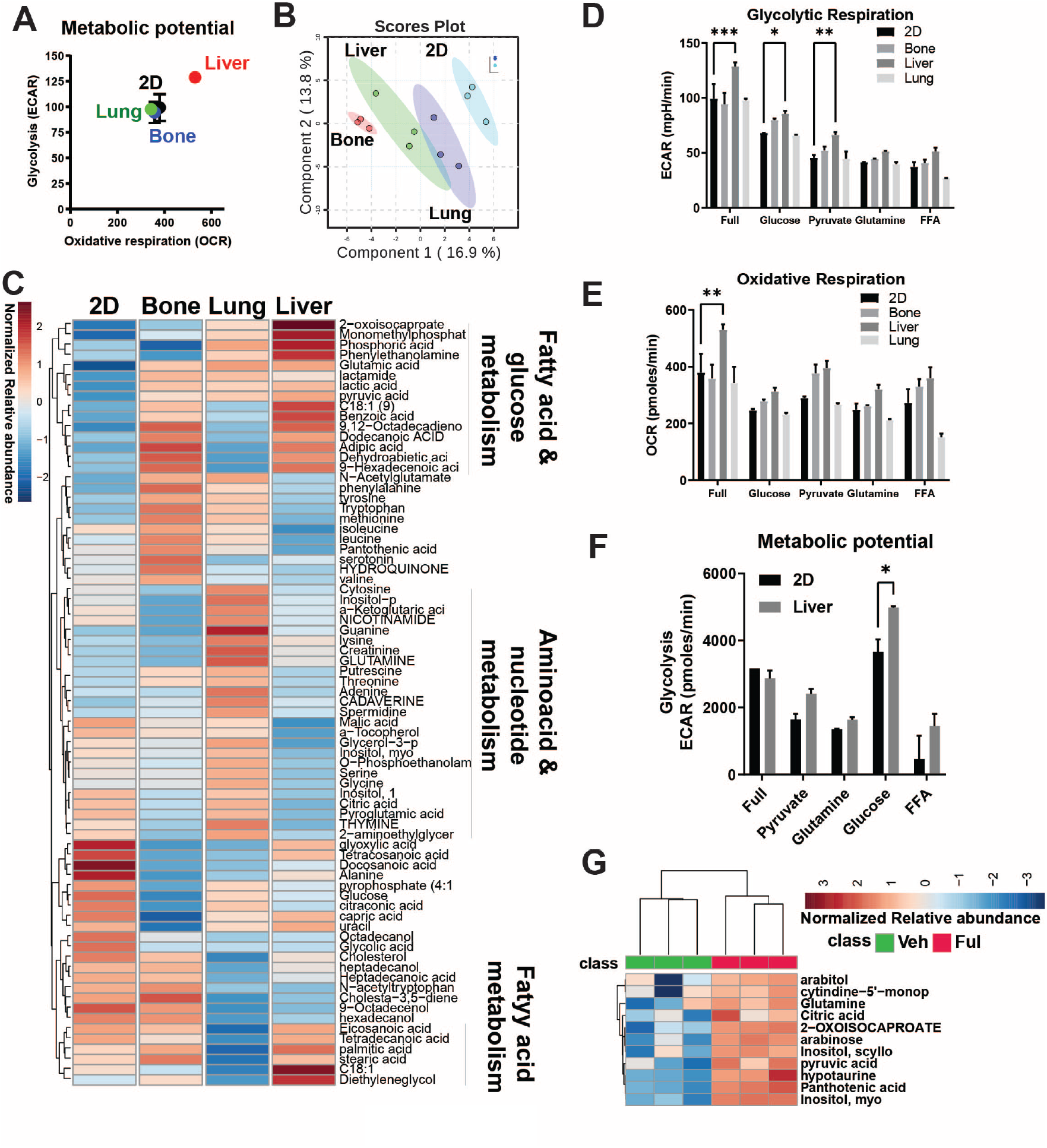
Glucose dependency is increased in MCF7-ESR1^Y537S^ cells cultured in liver ECM with Fulv treatment. **A)** Cell metabolic phenotype assay using the Seahorse Cell Energy Phenotype Kit. Cells cultured in plates coated with bone, liver, and lung Native Coat ECMs for 24 hours were tested for the energy phenotype. Each experiment was replicated twice with three technical replicates. Results from a representative experiment are shown. **B)** Whole metabolite profiling using GC/MS analysis of extracts from MCF7-ESR1^Y537S^ grown on plastic (Ctrl), bone, lung, and liver Native Coat ECMs. Scores plot showing distinct metabolite abundance patterns on different ECMs. **C)** Heatmap of metabolite profiling analysis of pathway changes induced by different ECMs. **D)** Cell phenotype assays were performed to dissect the specific fuel dependency on different ECMs, which showed that liver ECM but not bone or lung ECMs increased glycolytic respiration in the presence of full media, or media with glucose or pyruvate compared to cells that were grown on plastic. Two-way ANOVA, Dunn’s test, *p<0.05. **E)** Liver ECM led to an increase in oxidative respiration only when cells were supplemented with full media. **F)** Cell metabolic phenotype assay showed that MBC cells grown on liver ECM in media with glucose increased in glycolytic potential. **G)** Whole-metabolite profiling in MBC cells grown on plates coated with liver ECM and treated with vehicle (Veh) or 1 μM Fulv for 24 hours.

### Glucose metabolism pathways are upregulated in liver metastatic tumors in vivo

To uncover the mechanistic basis of decreased Fulv efficacy in patients with ER^+^ liver MBC, we used a preclinical xenograft mouse model with MCF7-ESR1^Y537S^ cells. Tail vein injection of MCF7-ESR1^Y537S^cells in NSG mice resulted in liver metastasis, with a small portion of tumors forming in the lung, and Fulv treatment failed to reduce metastatic burden (**Fig. 4A and 4B**). To study if metastatic site altered gene regulation in response to Fulv, we compared gene expression changes of liver and lung xenograft tumors. This analysis showed that Fulv reduced expression of classical ER target genes PgR and GREB1 in both sites in MCF7-ESR1^Y537S^ xenografts. However, unlike lung metastatic tumors, liver metastatic tumors failed to downregulate cell cycle genes, *MKI67* and *PCNA* in response to Fulv (**Fig. 4C**). Similar pattern of gene regulation was observed in MCF7-ESR1^D538G^ xenografts (**Figure S4A and S4B**). To validate this observation, we performed spatial sequencing in Veh and Fulv treated liver metastatic tumors from MCF7-ESR1^Y537S^ xenografts. Overall, UMAP plots showed that tumors from same treatment groups clustered together (**Fig. 4D**). We identified 17 distinct clusters associated with liver metastatic tumors (**Fig.4E**). Human transcripts were majority of the transcripts identified in liver tumor sections we analyzed (**Fig. 4F)**, while mouse specific transcripts for different cell types localized to tumor periphery **(Figure S4C**). Interestingly, Fulv treatment increased ESR1 mRNA expression throughout the tumors. While *MKI67* or *PCNA* expression did not change in response to Fulv, classical ER target changes *GREB1* and *PgR* were downregulated (**Fig. 4F**). When we analyzed enrichment of transcripts from cell populations with high MKI67 expression, cell cycle related genes were common to high MKI67 expression populations from both Veh and Fulv treated tumors. Of note, high MKI67 expressing populations from Fulv treated tumors had increased expression of gene sets associated with chromatin organization, carbohydrate metabolism and ribosome biogenesis (**Table S2**). These results showed that, while ERα retained some of its activity and Fulv response in liver metastatic tumors, suppression of cell proliferation in response to Fulv was lost (**Figure S5A**).

**Figure 4:**
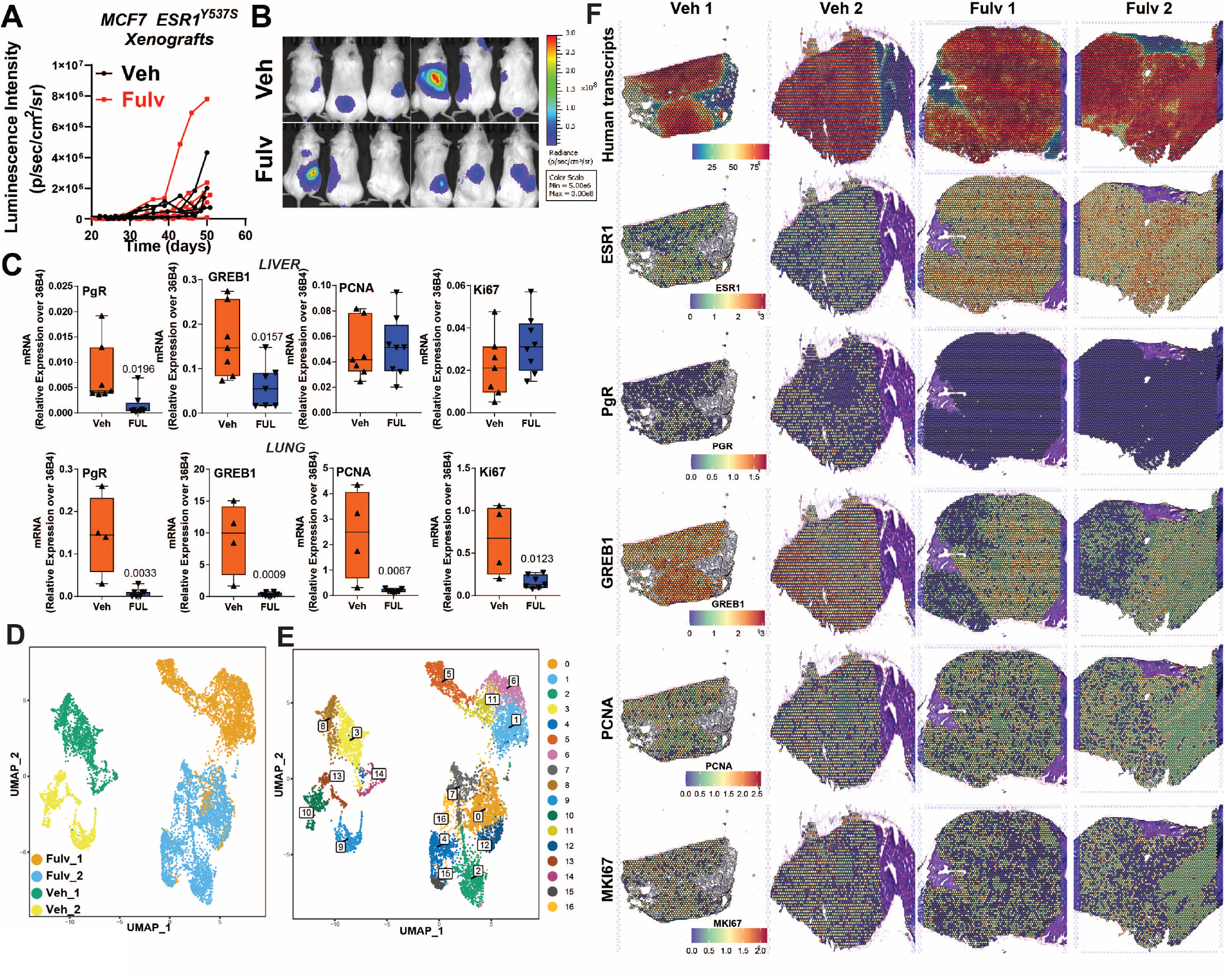
ER^+^ Liver metastatic xenograft tumors do not respond to Fulv. **A)** MCF7-ESR1^Y537S^ cells were grafted intravenously (tail vein), and mice were treated with placebo or Fulv. Metastatic burden was measured using IVIS to detect bioluminescence. N=6, metastatic tumor luciferase signal from each mouse is plotted. No statistically significant difference was detected in metastatic burden with Fulv using a two-way ANOVA for Fulv effect over time. **B)** Imaging of luciferase signal indicating metastatic outgrowth in the livers, lungs, and bones. **C)** RNA was isolated from the MCF7-ESR1^Y537S^ xenograft liver tumors. mRNA expression of ERα target genes GREB1 and PgR, and cell cycle-related genes PCNA and Ki67 were measured by qRT-PCR. Unpaired t-test, p values are indicated. **D)** UMAP plots of Visium spots from 2 Veh and 2 Fulv tumors. UMAP plots **E)** 17 clusters were identified **F)** Level of total and specific human transcripts identified in xenograft tumors

### Tumor metastatic burden correlates with carbohydrate levels in the diet

To determine why Fulv failed to reduce tumor progression and if any critical survival pathways were activated in response to Fulv, we performed RNASeq of liver metastatic tumors (**Fig. 5A**). Fulv significantly increased glycolysis and glycogenesis metabolism pathways, as well as lipid metabolism-related genes, consistent with our metabolomics analysis in **Fig. 3** (**Fig. 5B**). We compared changes in expression of genes in cells that were grown on plastic (2D) treated with Fulv, 4-hydroxytamoxifen, or palbociclib; different ECMs (Bone, liver, or lung); or changes in mouse transcripts in xenograft tumors with Fulv treatments. We found that glycogen pathway regulation was strongest in xenograft samples, suggesting that glycogen pathway regulation occurs in liver MBC, but not when cells are treated with Fulv in 2D or even when cells were grown on liver ECM without other components of the tumor microenvironment (**Fig. 5B**). Changes in glucose and lipid metabolism genes were observed in liver but not lung metastatic tumors (**Figure S5B**). Because we observed major metabolic and gene expression changes in glucose and lipid metabolism and an increased dependence on and utilization of glucose in models of liver metastasis, we next tested the impact of different diets with varying carbohydrate and fat contents in our MCF7-ESR1^Y537S^ xenograft model. Animals were provided with a Control diet, HFD, and FMD, which enabled us to examine impact of varying glucose or fat levels in the diet. Metastatic burden was measured over 4 weeks (**Fig. 5C and 5D**). At the end of the experiment, livers from animals who consumed the FMD looked healthier and had less visible liver metastasis nodules (**Fig. 5E**). Throughout the study we also monitored animal weight and caloric intake. Using this data, we performed a correlation analysis using all the data we obtained from these measurements and diet composition data, which revealed that metastatic burden is positively correlated with carbohydrate levels, particularly with di- and polysaccharides, in the diet. This finding is consistent with our observation that there was an increased dependence on glucose metabolism in liver metastatic tumors (**Fig. 3**). We found that metastatic burden negatively correlated with fat intake levels, while no correlation was observed for protein levels in the diet (**Table S3**). Interestingly, PAS staining for glycogen revealed that in animals who were fed a control diet or HFD, glycogen deposition was primarily in tumors (**Fig. 5F**).

**Figure 5:**
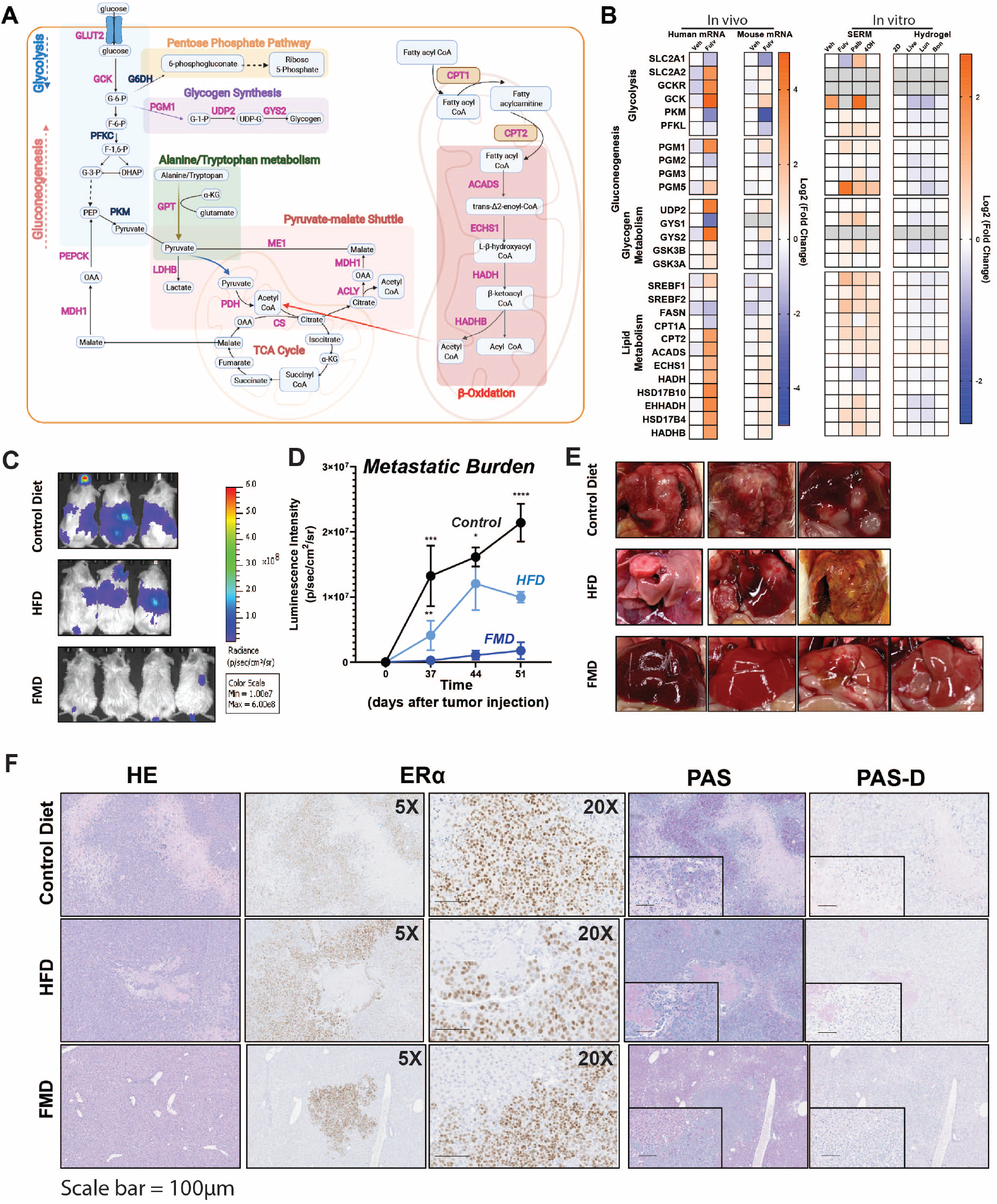
Metastatic burden increases with increasing carbohydrate percentage in the diet. **A)** RNA was isolated from the liver metastatic tumors of MCF7-ESR1^Y537S^ xenografts and RNASeq was performed using 6 samples from each treatment group. Fulv significantly increased glycolysis and glycogenesis metabolism pathways, as well as lipid metabolism. Pink-labeled genes are upregulated, and blue-labeled genes are downregulated. **B)** Comparison of glycolysis, glycogenesis, and lipid metabolism-related genes in the MCF7-ESR1^Y537S^ liver tumors, mouse transcripts identified in these tumors, in MCF7-ESR1^Y537S^ cells grown plastic re treated with Veh or 1μM Fulv, Palb or 4OHT, and MCF7-*ESR1*^Y537S^ cells grown on plates coated with different ECMs. Fold change relative to Veh-treated samples is plotted in each dataset. **C)** NSG mice with MCF7-ESR1^Y537S^ xenografts were fed a control diet, HFD, or FMD. Metastatic burden was measured using IVIS to detect bioluminescence. **D)** Mean Luminescence intensity of tumors was plotted to determine metastatic burden. N=3, Two-way ANOVA, Tukey’s post hoc test, * p <0.05, ***p <0.001, **** p <0.0001. Bars represent SEM. **E)** Livers from corresponding animals in C. **F)** Histological analysis of tumors from (A). HE, ERα, PAS, and PAS-D IHC staining were performed.

### Targeting glycogen deposition using dietary approaches improve Fulv response of liver metastatic tumors

Because we observed major metabolic and gene expression changes in glucose and glycogen metabolism genes, and an increased dependence on and utilization of glucose in models of liver metastasis, we next tested the impact of an FMD on MCF7-ESR1^Y537S^ xenograft model. FMD prevents glycogen accumulation in the liver and thus blocks a glucose surge and resultant insulin release from the pancreas. In addition, a recent study combining FMD with Fulv showed a reduced time to endocrine resistance in primary ER^+^/HER2^-^ mouse tumors and achieved complete response or stable disease in human patients (56). To determine whether reducing glucose metabolism using an FMD might synergize with Fulv treatment in our in vivo liver metastasis model, we delivered MCF7-ESR1^Y537S^ cells through tail vein injection in NSG mice. Consuming FMD synergized with Fulv treatment to reduce metastatic burden in mice with metastatic tumors (**Fig. 6A and 6B**) and decreased the number of visible metastatic nodules in the liver (**Fig. 6C and 6D**). Fulv treatment increased glycogen deposition in tumors (**Fig. 6E, column 3, and row 2**). These data provide proof of concept that combining a dietary intervention targeting glucose metabolism and glycogen deposition in liver provided a durable response to Fulv.

**Figure 6:**
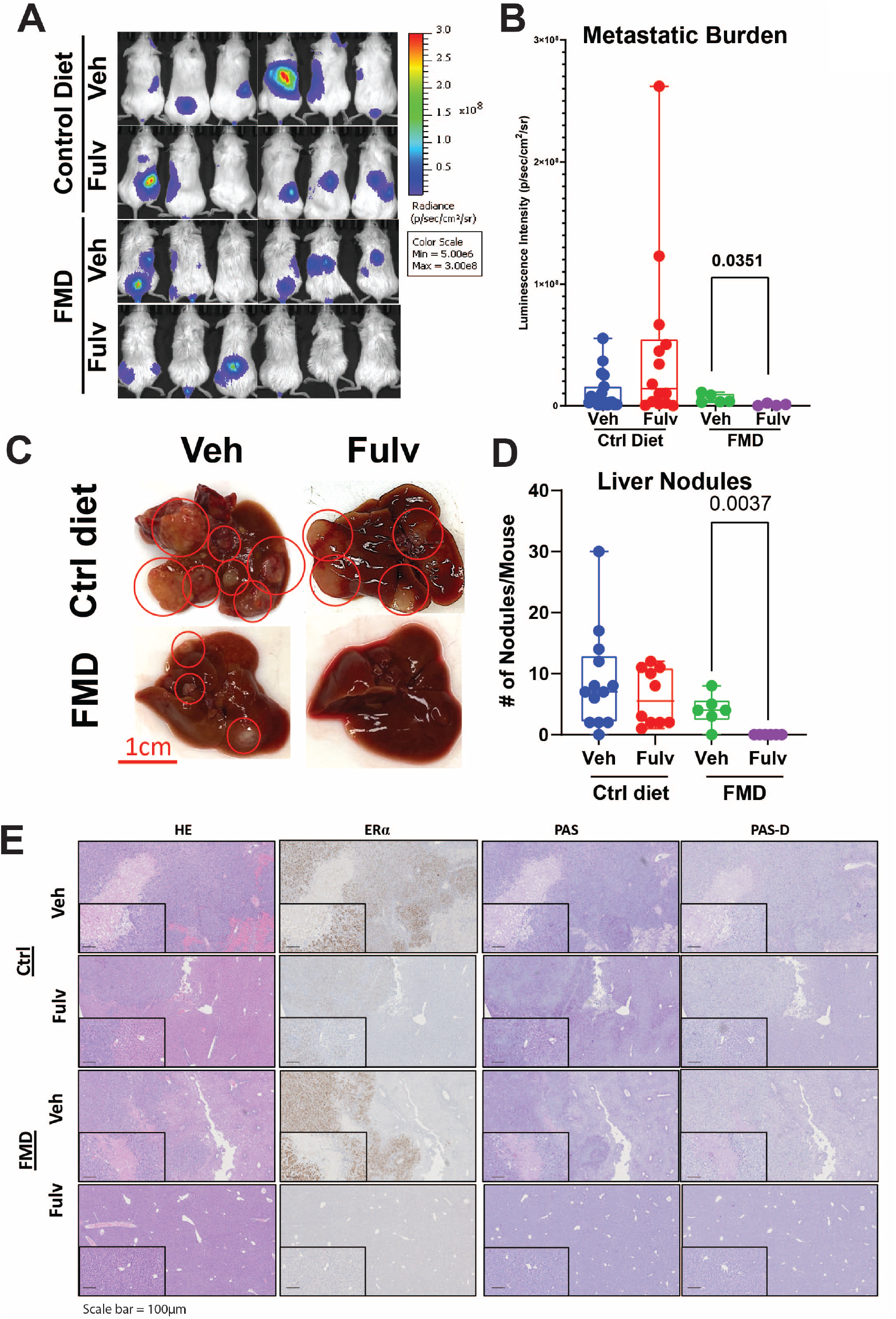
FMD diet synergizes with Fulv to reduce MCF7-ESR1^Y537S^ liver metastatic burden and number of nodules. **A, B)** NSG mice with MCF7-ESR1^Y537S^ xenografts were fed control diet or FMD. Mice were treated with Veh or Fulv. Bioluminescence imaging of tumors was measured by IVIS. (Image for the control group is same as the image for Figure 4B) **C)** Representative images of livers from mice in different treatment groups. **D)** Liver metastatic nodules were counted at necropsy. One-way ANOVA, Dunn’s multiple comparison test, *p<0.05. **E)** Histological analysis of tumors from (C). HE, ERα, PAS, and PAS-D IHC staining were performed.

## Discussion

Metastatic tumor phenotypes and responses to treatment reflect key aspects of the tumor microenvironment (57). Here, we investigated tumor-intrinsic metabolic mechanisms that arise specifically in the liver metastatic niche. Using a combination of models, spatial and molecular data, and analytical methods, our findings reveal a key mechanism of endocrine resistance in liver MBC: niche-related metabolic plasticity in MBC cells that alters the response to ER-targeted therapies. This metastatic niche displays specific metabolic changes that provide a mechanistic basis for the poor performance of Fulv treatment on survival of patients with ER^+^ liver MBC. Yet, these changes also unveiled a unique metabolic vulnerability that could be exploited through dietary interventions to improve Fulv response.

Treatment of MBC in the clinic focuses on targeting mechanisms that underlie increased activity of selective cellular pathways. Indeed, interrogating and targeting signaling cascades in tumors has been fruitful in promoting the development of endocrine-based therapies combining CDK4/6 inhibitors, PI3K inhibitors, or mTOR pathway inhibitors with endocrine agents for ER^+^ MBC (13). Yet, none of these therapies is metastatic site-specific, and tumors can still develop resistance to combination therapies. In such cases, the cancer that develops is considerably more aggressive due to hyperactivation of compensatory pathways (58). In contrast, we focused on inhibiting dynamic metabolic mechanisms enabling survival and therapy resistance in liver metastatic niches. Targeting metastatic site-specific metabolic vulnerabilities in ER^+^ MBC is a novel approach and the impact of metabolic interventions on endocrine therapy effectiveness is underexplored.

To characterize ER antagonist responses of MBC cells and test potential efficacy of combination therapies, we previously used decellularized tissue-specific ECM hydrogels from different metastatic tissues (19). Despite lacking non-tumor cells in tumor microenvironment, hydrogels constitute compatible niches to support ER^+^ MBC growth and provide an opportunity to recapitulate metastatic site environment and analyze drug responses and metastasis-associated phenotypes in vitro. Characterization of hydrogel composition by mass spectrometry and quantitative biochemical and biophysical assays showed that hydrogels retain tissue-characteristic ECM proteins as well as growth factors (19,59,60). Scanning electron microscopy analysis revealed structural similarity to human tissues, and rheometry analysis showed conservation of biophysical properties of the hydrogels, such as stiffness and resistance to deformation, as well as response of cancer cells grown in these matrices to drugs (19,61). We found that, consistent with the clinical observation, breast cancer cells grown on liver hydrogels were less responsive to Fulv compared to cells grown on Matrigel or lung or bone hydrogels. Supporting our observations in the hydrogel systems, an in vitro study reported reduced Fulv response when MCF7-ESR1^Y537S^ were cultured with human hepatocytes (62).

ESR1 activating mutations are detected in metastatic tumors from patients who were previously treated with aromatase inhibitors (47-50). These mutations are reported to be enriched in liver metastatic tumors (51-53). Using two publicly available datasets (54,55) and data from our clinical cohort, we validated enrichment of ESR1 mutations in liver metastatic tumors. In our cohort, patients with liver metastasis had worst survival independent of the ESR1 status. Even though we mainly used ESR1 mutant cell models to ensure modeling metastatic disease, cell lines with wild type ERα also displayed similar response to Fulv in liver hydrogels. Our findings do not rule out a larger impact of ESR1 mutations on metastatic disease outcomes, however our observations support a more general regulatory role for liver microenvironment in rendering Fulv less efficacious in ER^+^ MBC cells independent of the ESR1 mutation status.

We observed altered ERα activity and recruitment in different tissue-specific ECMs. Since ERα expression is the same in different hydrogels, the altered ER recruitment pattern likely does not result from altered ER expression. While MBC grown in bone ECM have increased ER signaling, those MBC grown in liver ECM had fewer ER binding sites that associate with liver ECM modulated genes. Epigenetic marks are well established as a mechanism that dictates ERα recruitment to chromatin, target gene regulation, and response to ER antagonists (63-65). Histone 3 displays altered acetylation at lysine 4, 9, and 27 in breast cancer cells that fail to respond to ER antagonists (65-67). Systemic metabolic changes impact hormone-dependent cancer cell metabolism (41,68) and composition of histone marks (39,69), and response to therapies (19,41). Our findings suggest an altered ERα activity in MBC cells residing in the liver microenvironment, and a role for metabolic and epigenetic enzymes to interact with ER in a Fulv-dependent manner to locally change epigenetic marks, which would lead to altered response to ERα antagonists. Additionally, metastatic site adaptation (70,71) and metabolic alterations in MBC cells result in epigenetic reprogramming due to changes in the availability of substrates for epigenetic enzymes (72-74). Local acetyl-CoA production via recruitment of metabolic enzymes to chromatin enables coordination of environmental cues with histone acetylation and gene transcription, thus increasing fitness and survival of cancer cells in different metastatic tissues. Acetyl-CoA can be synthesized in the nucleus from pyruvate by the pyruvate dehydrogenase complex (PDC) (75,76), from acetate by acetyl-CoA synthetase 2 (ACSS2) (77,78), or from citrate by ATP-citrate lyase (ACLY) (79). Intriguingly, all these enzymes are upregulated in our ER^+^ liver metastatic xenografts upon Fulv treatment, supporting a role for altered metabolism in changing tumors’ epigenetic landscape. Lack of major changes in glycolytic and mitochondrial respiration of MCB cells in response to Fulv in liver ECM support utilization of products of these pathways for other regulatory pathways. Future studies are needed to uncover how increased glucose utilization change substrates of epigenetic enzymes and how these metabolites result in epigenetic differences based on metastatic sites.

Accumulating data indicate that practical clinical dietary intervention during cancer treatment has a profound impact in improving the efficacy of anticancer therapy, especially the FMD (80). This diet is generally high-fat, moderate-protein, and very-low-carbohydrate, leading to a process known as ketogenesis (81). Lipolysis-induced fatty acids are metabolized to acetoacetate, which is later converted to β-hydroxybutyrate (β-OHB) and acetone (82). β-OHB is the most abundant ketone body derived from β-oxidation in the liver, and it replaces glucose as a primary source of energy (82). Considering the impact of carbohydrates in promoting breast cancer cell proliferation, the FMD has the potential to limit or control tumor growth. Some animal studies support that an FMD inhibits not only the progression of the primary tumor but also systemic metastasis (83,84). By providing a low-glucose microenvironment, the FMD enhances the cancer cell therapeutic response through selective metabolic oxidative stress (85). Recent evidence indicates that the combination of FMD with Fulv results in increased time to endocrine resistance in primary ER^+^/HER2^-^ mouse tumors and in complete response or stable disease in human patients (56). This important study focused on the impact of FMD on circulating factors and signaling pathways; we instead focus on the impact of FMD on metabolic state of tumor cells. Recently, suppression of insulin feedback after PI3K inhibition using a FMD was proposed (86). Our findings further support use of this diet to improve endocrine therapy responses for MBC patients with liver metastasis.

We present a wealth of data from a clinical cohort of metastatic breast cancer patients, in vitro 3D models and in vivo xenografts to support a role in liver microenvironment in changing response of breast cancer cells to Fulv. One of the limitations of our study is the use of immunocompromised mouse models that lack certain immune cell populations. Immune system is an essential regulator of metastatic growth and therapeutic resistance. Future studies are needed to develop liver metastasis specific syngeneic mouse models to dissect impact of Fulv and diet in the context of a fully functional immune system. Use of liver metastatic tumor biopsy samples might enable study of gene expression and metabolic phenotypes in human patients. Co-culturing of stromal and immune cells together with MBC in 3D models might improve utility of hydrogel systems to elucidate the role of tumor microenvironment on metabolic and epigenetic changes that lead to reduced Fulv efficacy. Finally, analysis of more current data from clinical trials that incorporate new combination strategies, including CDK4/6 inhibitors, PI3K inhibitors, mTOR inhibitors, would help improve our understanding of how these more contemporary strategies impact disease outcome for patients with liver MBC.

In summary, our studies established metastatic-niche specific metabolic vulnerabilities as a novel target by uncovering the potential of FMD to improve Fulv response in ER^+^ liver MBC. We envision that metabolism-based therapies in combination with standard endocrine-based therapies may be effective in exploiting metastatic site-specific metabolic dependencies of cancer cells, and in eliciting durable responses. Given the need for better strategies to treat liver metastatic tumors, our work offers both novel metabolic insights and a more complete understanding of the basic molecular mechanisms that underlie drug resistance. This novel understanding will enable us and others to exploit these new vulnerabilities to improve therapy response of MBCs, and reduce morbidity and mortality associated with liver metastasis.

## Acknowledgements

Clinical data were provided from the Breast Cancer Management System database that is maintained and managed by the Department of Breast Medical Oncology from the University of Texas MD Anderson Cancer Center. We would like to thank members of UIUC Roy J. Carver Biotechnology Center. Specifically, Lucas Li and Alex Ulanov for metabolomics analysis and Alvaro Hernandez, Chris Wright and Kate Janssen for bulk and spatial sequencing analysis. We would like to thanks Karen Doty for assistance with histology analysis. This work was supported by grants from the University of Illinois, Office of the Vice Chancellor for Research, Future Interdisciplinary Research Endeavors grant from college of ACES, a seed grant from Cancer Center at Illinois (to ZME), and National Institute of Food and Agriculture, U.S. Department of Agriculture, award ILLU-698-909 (to ZME). Research reported in this publication was supported by the Cancer Scholars for Translational and Applied Research (CSTAR) Program sponsored by the Cancer Center at Illinois and the Carle Cancer Center under Award Number CST EP082021 (to ANM). The content is solely the responsibility of the authors and does not necessarily represent the official views of the program sponsors. Research reported in this publication was supported by the National Institute of Biomedical Imaging and Bioengineering of the National Institutes of Health under Award Number T32EB019944 (to ASC). The content is solely the responsibility of the authors and does not necessarily represent the official views of the National Institutes of Health. BHP is supported by the Susan G. Komen Foundation and the Breast Cancer Research Foundation. BSK was supported by a Breast Cancer Research grant 20-083.

