## Supplementary material for "Targeting metabolic adaptations in the breast cancer–liver metastatic niche using dietary approaches to improve endocrine therapy efficacy": Table S3

| | P (two-tailed) | Pearson r | R squared | P value summary |
|---|---|---|---|---|
| Calories(kcal/r) | 0.0492 | -0.997 | 0.994 | * |
| Caloric Intake (kcal/Body weight) | 0.946 | -0.08474 | 0.007181 | ns |
| Body weight | 0.213 | 0.9445 | 0.8922 | ns |
| % Carbohydrate | 0.0244 | 0.9993 | 0.9985 | * |
| % Fat | 0.0103 | -0.9999 | 0.9997 | * |
| % Protein | 0.2675 | 0.913 | 0.8336 | ns |
| Carb. cal (kcal/g) | 0.0233 | 0.9993 | 0.9987 | * |
| Fat (kcal/g) | 0.0108 | -0.9999 | 0.9997 | * |
| Prot. (kcal/g) | 0.2675 | 0.913 | 0.8336 | ns |
| Monosaccharides | 0.2039 | -0.9492 | 0.9009 | ns |
| Disaccharides | 0.0253 | 0.9992 | 0.9984 | * |
| Polysaccharides | 0.0339 | 0.9986 | 0.9972 | * |
| 18:2 Linoleic acid | 0.097 | -0.9884 | 0.9769 | ns |
| 18:3 Linoleic acid | 0.0536 | -0.9965 | 0.9929 | ns |
| Total saturated fat | 0.0022 | -1 | 1 | ** |
| Total monounsaturated fat | 0.0716 | -0.9937 | 0.9874 | ns |
| Total polyunsaturated fat | 0.0899 | -0.99 | 0.9802 | ns |
| Ala | 0.2675 | 0.913 | 0.8336 | ns |
| Arg | 0.2675 | 0.913 | 0.8336 | ns |
| Asp | 0.2675 | 0.913 | 0.8336 | ns |
| Cys | 0.2675 | 0.913 | 0.8336 | ns |
| Glutamate | 0.2675 | 0.913 | 0.8336 | ns |
| Glycine | 0.2675 | 0.913 | 0.8336 | ns |
| Hist | 0.2675 | 0.913 | 0.8336 | ns |
| Iso | 0.2675 | 0.913 | 0.8336 | ns |
| Leu | 0.2675 | 0.913 | 0.8336 | ns |
| Lys | 0.2675 | 0.913 | 0.8336 | ns |
| Methionine | 0.2675 | 0.913 | 0.8336 | ns |
| Phen | 0.2675 | 0.913 | 0.8336 | ns |
| Prot cal | 0.2675 | 0.913 | 0.8336 | ns |
| Ser | 0.2675 | 0.913 | 0.8336 | ns |
| Thr | 0.2675 | 0.913 | 0.8336 | ns |
| Tryp | 0.2675 | 0.913 | 0.8336 | ns |
| Tyr | 0.2675 | 0.913 | 0.8336 | ns |
| Val | 0.2675 | 0.913 | 0.8336 | ns |
