## Supplementary figures and images for "Targeting metabolic adaptations in the breast cancer–liver metastatic niche using dietary approaches to improve endocrine therapy efficacy"

### Figure S2

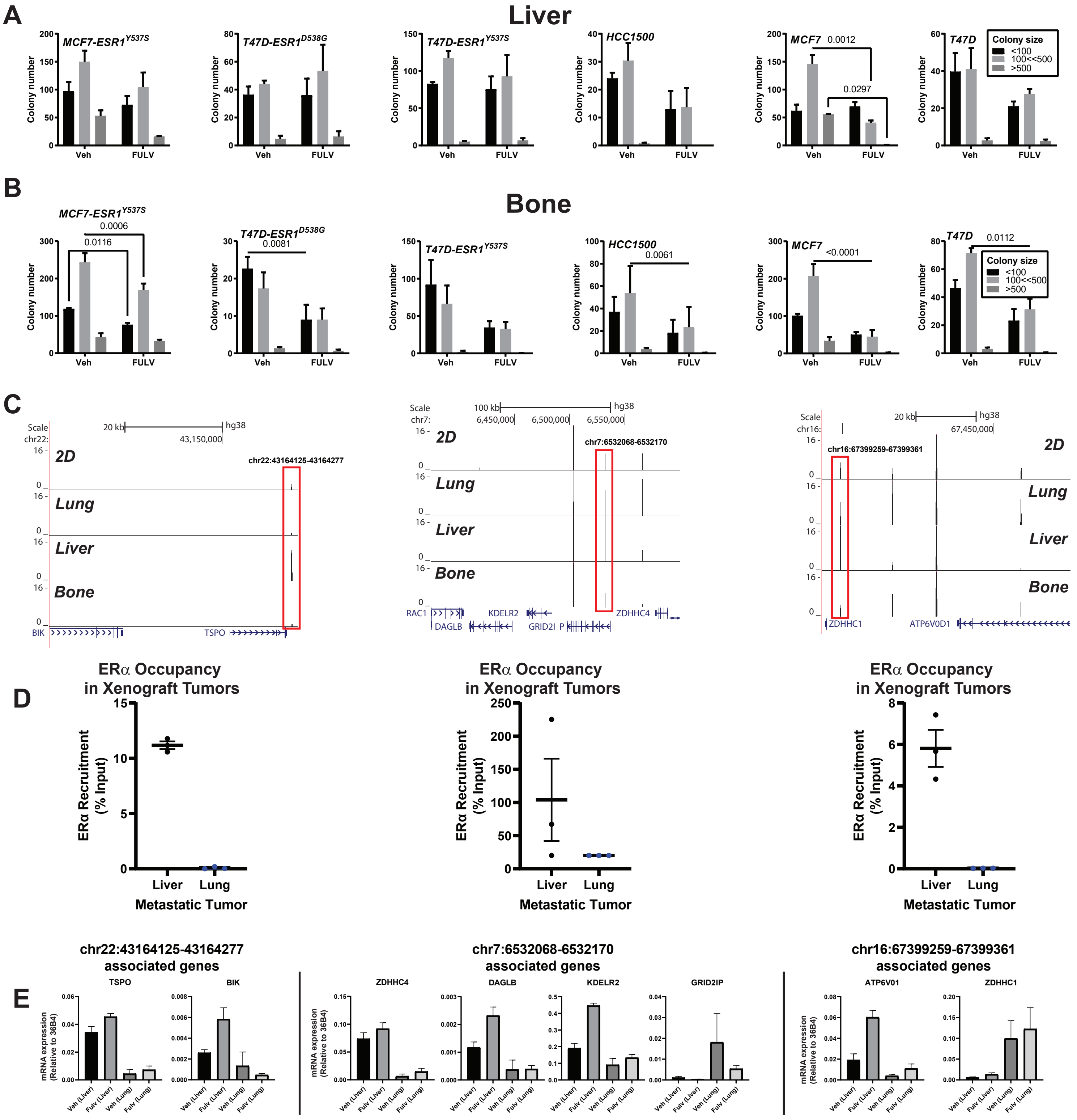


**Supplementary Figure 2**

### Figure S3

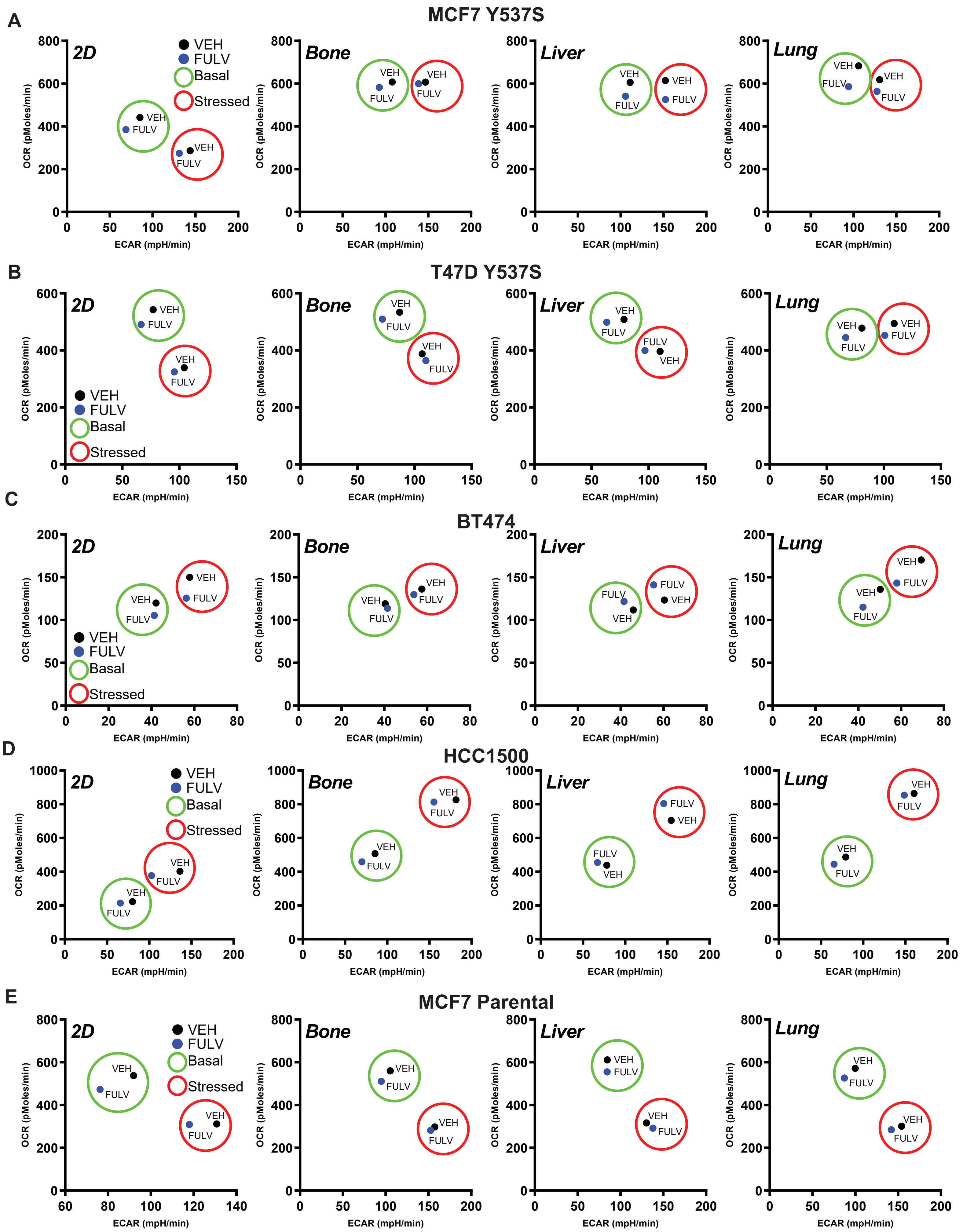


**Supplementary Figure 3**

### Figure S4

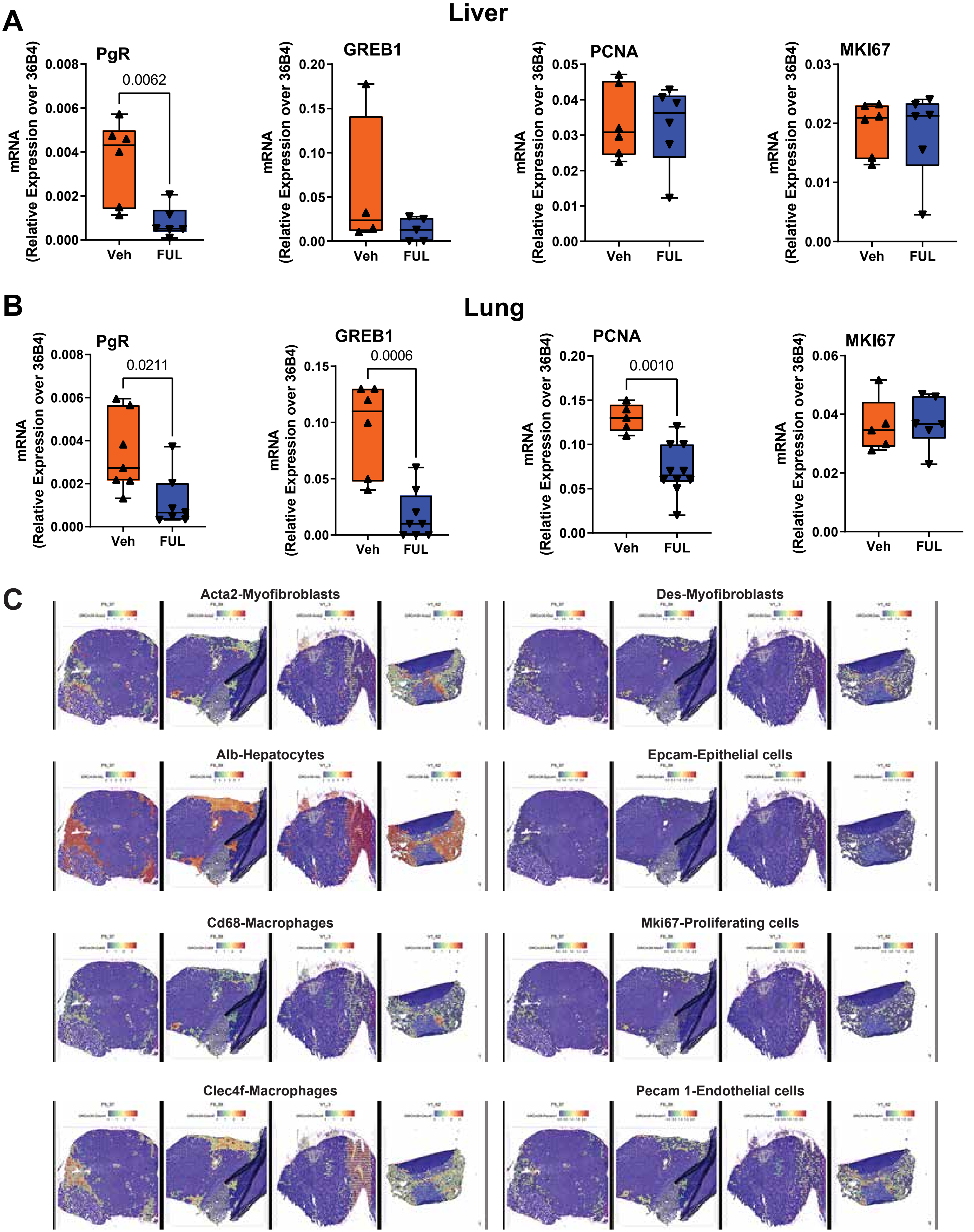


Supplementary Figure 4

### Figure S5

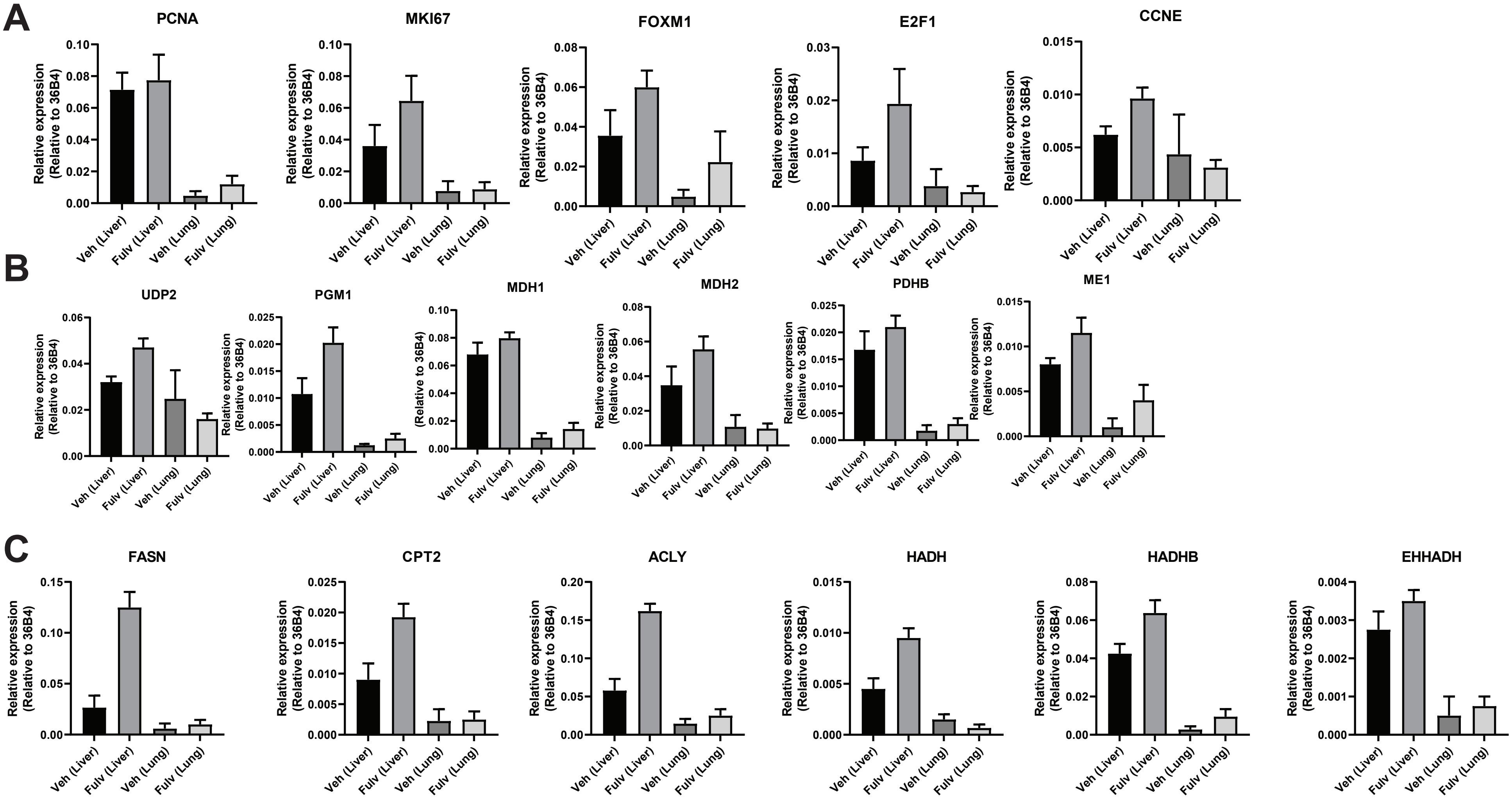


**Supplementary Figure 5**
