## Supplementary material for "Targeting metabolic adaptations in the breast cancer–liver metastatic niche using dietary approaches to improve endocrine therapy efficacy": Figure S1

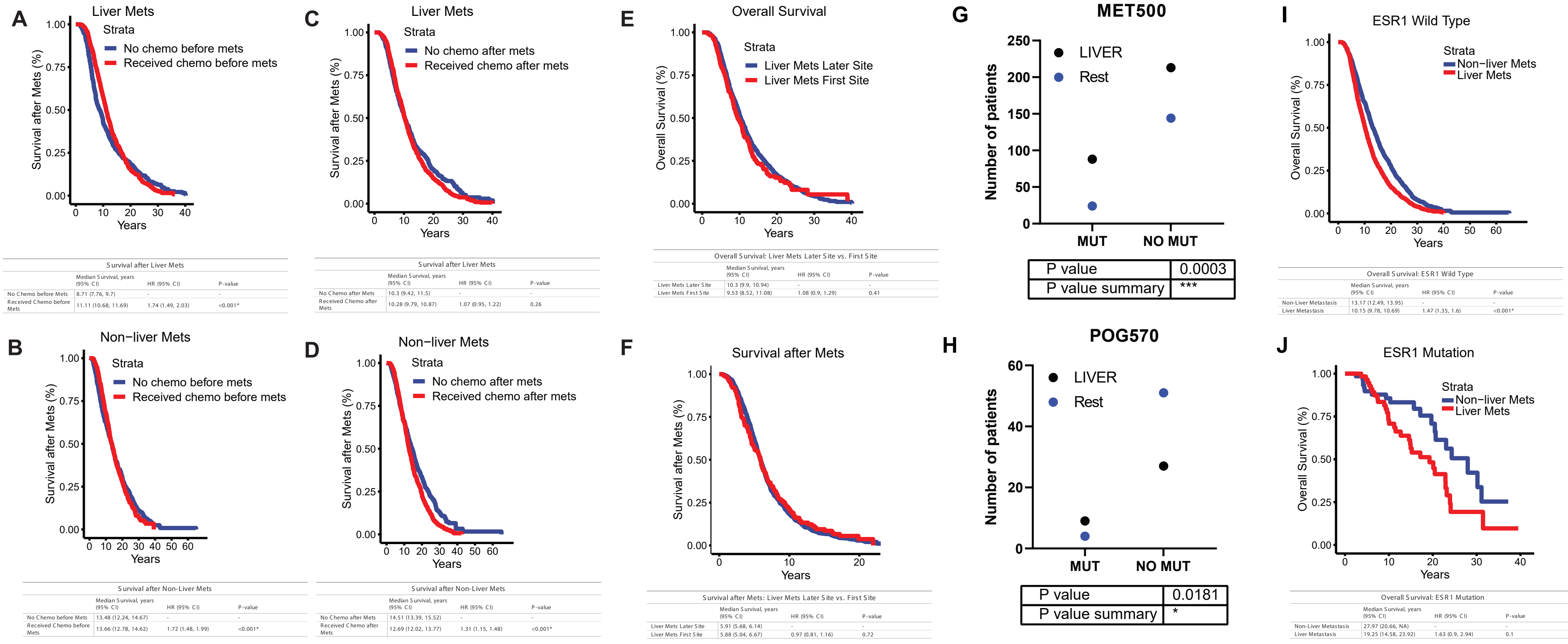


Survival after Liver Mets

| | Median Survival, years (95% CI) | HR (95% CI) | P-value |
|---|---|---|---|
| No Chemo before Mets | 8.71 (7.76, 9.7) | - | - |
| Received Chemo before Mets | 11.11 (10.68, 11.69) | 1.74 (1.49, 2.03) | <0.001* |

Survival after Liver Mets

| | Median Survival, years (95% CI) | HR (95% CI) | P-value |
|---|---|---|---|
| No Chemo after Mets | 10.3 (9.42, 11.5) | - | - |
| Received Chemo after Mets | 10.28 (9.79, 10.87) | 1.07 (0.95, 1.22) | 0.26 |

Overall Survival: Liver Mets Later Site vs. First Site

| | Median Survival, years (95% CI) | HR (95% CI) | P-value |
|---|---|---|---|
| Liver Mets Later Site | 10.3 (9.9, 10.94) | - | - |
| Liver Mets First Site | 9.53 (8.52, 11.08) | 1.08 (0.9, 1.29) | 0.41 |

Overall Survival: ESR1 Wild Type

| | Median Survival, years (95% CI) | HR (95% CI) | P-value |
|---|---|---|---|
| Non-Liver Metastasis | 13.17 (12.49, 13.95) | - | - |
| Liver Metastasis | 10.15 (9.78, 10.69) | 1.47 (1.35, 1.6) | <0.001* |

Survival after Non-Liver Mets

| | Median Survival, years (95% CI) | HR (95% CI) | P-value |
|---|---|---|---|
| No Chemo before Mets | 13.48 (12.24, 14.67) | - | - |
| Received Chemo before Mets | 13.66 (12.78, 14.62) | 1.72 (1.48, 1.99) | <0.001* |

Survival after Non-Liver Mets

| | Median Survival, years (95% CI) | HR (95% CI) | P-value |
|---|---|---|---|
| No Chemo after Mets | 14.51 (13.39, 15.52) | - | - |
| Received Chemo after Mets | 12.69 (12.02, 13.77) | 1.31 (1.15, 1.48) | <0.001* |

Survival after Mets: Liver Mets Later Site vs. First Site

| | Median Survival, years (95% CI) | HR (95% CI) | P-value |
|---|---|---|---|
| Liver Mets Later Site | 5.91 (5.68, 6.14) | - | - |
| Liver Mets First Site | 5.88 (5.04, 6.67) | 0.97 (0.81, 1.16) | 0.72 |

Overall Survival: ESR1 Mutation

| | Median Survival, years (95% CI) | HR (95% CI) | P-value |
|---|---|---|---|
| Non-Liver Metastasis | 27.97 (20.66, NA) | - | - |
| Liver Metastasis | 19.25 (14.58, 23.92) | 1.63 (0.9, 2.94) | 0.1 |

### Supplementary Figure 1
